# Bone marrow by day, lymph nodes by night: Undernutrition imparts a distinct circadian rhythm to T cells in mice

**DOI:** 10.1101/2024.05.24.595677

**Authors:** Takesha R. Foster, Kwesi A. Dadzie, Sydney Dunn, Melanie R. Gubbels Bupp

**Affiliations:** Department of Biology, Randolph-Macon College, Ashland, VA, United States

## Abstract

In well-nourished organisms, T cells migrate between the blood and secondary lymphoid organs, conducting surveillance for invading pathogens. T cell surveillance is under circadian control via diurnal fluctuations in corticosterone levels and undernutrition is associated with increased corticosterone. Therefore, we hypothesized that undernutrition disrupts the circadian migratory patterns of T cells. We report that compared to well-nourished controls, undernourished mice demonstrate enhanced T cell relocation to the bone marrow throughout the 24-hour period, but especially during the light phase, and diminished T cell migration to the lymph nodes only during the light phase. Undernutrition-related changes in T cell expression of key migration proteins are also mostly limited to the light phase. For example, undernourished naïve CD4+ T cells exhibited higher levels of CXCR4 and CCR7 as well as reduced levels of S1P1 compared to controls; with all changes, except for CXCR4 expression, being restricted to the light phase. These results suggest that naïve CD4+ T cells in the lymph nodes upregulate CXCR4 during the dark phase, enabling their migration to the bone marrow where they remain for the light phase. Once there, CCR7 is upregulated, presumably sending them back to the lymph node, thereby preserving immunosurveillance during the dark phase. Naïve CD4+ T cell disengagement from S1P1-related egress signals may further contribute to increased retention of cells within each compartment during the appropriate phase. Undernutrition-related increases in T cell residency of the bone marrow likely preserve T cell numbers until nutrition is restored.

## Introduction

Organisms face a trade-off between immunity, growth, and reproduction at all times, but especially during times of undernutrition [1]. As food intake decreases, animals invest fewer resources in their immune system, diminishing the ability of the host to resist invading pathogens [1–6]. Indeed, immunodeficiency is a hallmark of undernutrition and, in 2021, undernutrition was associated with 45% of deaths in children under the age of 5 due to increased risk and severity of acute infections, including diarrhea and acute respiratory illnesses [7–9]. Further, some vaccination efforts are less effective in undernourished children, due, at least in part, to reduced T cell functionality [7]. T cells isolated from undernourished children exhibit reduced proliferative capability and secrete less IL-2 and IFN-**γ** in response to stimulation than controls [10,11].

Circadian rhythms enable organisms to anticipate and respond to daily changes in their environment. They are established by the specialized suprachiasmatic nucleus (SCN) in the hypothalamus, which receives the zeitgeber, light, to entrain its phase [12]. Nearly all cells have internal “clocks” as well, many of which are entrained to some degree, by feeding rhythms [13]. Naïve and memory T cell surveillance of lymphoid organs is essential for timely adaptive immune responses [14]. Circadian control of T cell surveillance potentially increases the chance of a T cell encountering its cognate antigen, by positioning both T cells and antigen-presenting cells in lymphoid organs during the active period [14–16]. Specifically, T cells are primarily located in the blood during the inactive period, while peak T cell residency in the lymph nodes and spleen occurs during the active period, when the chances of encountering a pathogen are highest [16].

Glucocorticoids (GCs) are steroid hormones released by the adrenal cortex in a circadian fashion and regulate many biological processes, including the circadian nature of T cell surveillance [15,17]. In nocturnal animals like mice, blood GC levels peak at the start of the dark phase and fall during the light phase [12,15,16,18]. T-cell specific deficiency of the glucocorticoid receptor disrupts the diurnal distribution of T cells, revealing a critical role for GCs in the circadian nature of T cell surveillance [16]. Additional experiments revealed the underlying pathway: GCs regulate the diurnal expression of the IL-7 receptor alpha (CD127), which in turn controls the diurnal expression of CXCR4, a chemokine receptor for stromal-derived-factor-1 (SDF-1, also known as CXCL12) [16]. CXCL12 is expressed in the bone marrow and is known to be important for the homing, retention, survival, and quiescence of hematopoietic stem cells [19]. In well-nourished mice, circadian control of CXCR4 expression is associated with homing naïve and memory CD4+ T cells to the spleen during the dark phase, while during dietary restriction, CXCR4 is instead associated with CD8+ memory T cell homing to the bone marrow [16,20]. The expression of other key molecules in adhesion and homing of T cells also oscillate in a circadian fashion, including the chemokine receptor, CCR7, and S1P1 [21,22], which are associated with T cell homing and egress from the lymph nodes during the dark and light phases, respectively [22]. The egress function of S1P1 can be downregulated by CD69, an early activation marker [23], which also plays a role in effector CD4+ T cell migration to, or persistence in, the bone marrow [24,25]. Importantly, immune responses initiated in mice with intact diurnal T cell surveillance are functionally superior to those initiated in mice with disrupted diurnal T cell surveillance [16].

We and others have previously reported elevated levels of GC and heightened expression of CD127 on T cells during undernutrition [20,26]. Given the importance of GC and CD127 on the circadian rhythm [12,16,18], we hypothesized that the circadian rhythms of T cells may be disrupted in undernourished mice, perhaps contributing to diminished immunity. Therefore, we determined the total number of naïve and memory CD4+ and CD8+ T cells in the spleen, blood, lymph nodes, and bone marrow at regular intervals throughout a 24-hour period in under- and well-nourished mice. We also examined the expression levels of key molecules involved in T cell migration. Our findings reveal that naïve T cells in undernourished mice exhibit a distinct circadian rhythm, with increased migration to the bone marrow and diminished migration to lymph nodes during the inactive period.

## Materials and Methods

### Mice

All mice were housed in the mouse facility located at Randolph-Macon College. Male C57BL/6J mice were purchased from the Jackson Laboratory (Maine, USA) or bred in-house. All treatments were in accordance with the Institutional Animal Care and Use Committee guidelines (approved protocol number 21-02) and the National Institute of Health guide for the care and use of laboratory animals. C57BL/6J mice were singly housed, and chow (Teklad Global 18% Protein Rodent Diet, Harlan Laboratories) intake was monitored for two weeks. Mice randomly assigned to the undernourished (UN) treatment group received the lesser of either 35% less Teklad chow by weight than consumed two weeks prior or 13 grams. *Ad libitum* (AL) control mice received unrestricted access to 100g of Teklad chow. Both groups of mice received chow at approximately ZT12, the beginning of the dark phase. In all experiments, undernourishment lasted for one week and study mice were euthanized at 10 weeks of age via CO_2_ overdose followed by cervical dislocation. The overall reduction in average chow intake for undernourished mice was 56%. Although many individual undernourished mice lost up to 25% of their body weight, on average, undernourished mice lost 15%±1.5 while control mice gained 3%±0.5 of their original body weight in one week. This undernourishment protocol is similar to dietary restriction models that restrict food access by 50% but is distinct from calorie restriction [20,27].

### Corticosterone Enzyme Linked Immunosorbent Assay (ELISA)

Blood was extracted via cardiac puncture from male C57BL/6J mice at ZT0 and ZT12. Blood samples were allowed to clot for 30 minutes at room temperature and then centrifuged at 13,300 RPM for 20 minutes. Supernatants were collected and then stored at −80°C until ready for assaying. Corticosterone levels in serum diluted 1:40 were quantified using a corticosterone ELISA kit (ENZO Life Sciences, East Farmingdale, NY).

### Flow cytometry

Following the 1-week period of undernourishment or control treatment, cells were isolated from the spleens, blood, and bone marrow (both femurs) of male C57BL/6J mice. Cells isolated from the brachial, axillary, and inguinal lymph nodes were pooled. Red blood cells were cleared from the spleen, blood, and bone marrow preparations using ACK lysis buffer (0.15 M NH_4_Cl; 10 mM KHCO_3_; 0.1 mM EDTA). Following the lysis of red blood cells, 1.25 million cells were isolated from all tissue types and incubated with an Fc-shielding reagent (Purified Anti-Mouse CD16/CD32 (2.4G2)) (Cytek Biosciences, California, USA). Surface protein markers used to distinguish T cells of interest are as follows: leukocyte (CD45.2 (104, Cytek Biosciences)), naïve (CD62L (MEL-4, Biolegend, California, USA) and/or CD44 (IM7, Cytek Biosciences)), CD4+ T cell (CD4 (GK1.5, Thermofisher, Massachusetts, USA)), CD8+ T cell (CD8 (53-6.7, BD Biosciences, New Jersey, USA)). In addition, T cells were also stained for CXCR4 (2B11, eBioscience, San Diego, USA), CD127 (A7R34, Cytek Biosciences), CD69 (H1.2F3, Cytek Biosciences), S1P1 (713412, R&D Systems, Minneapolis, USA), and CCR7 (4B12, eBioscience). After staining, cells were washed and acquired on a NovoCyte 3005 flow cytometer (Agilent Technologies, California, USA). Data were analyzed using NovoExpress or FlowJo software.

### Statistical Analyses

Statistical analyses were conducted using GraphPad Prism. The statistical significance of differences between treatment groups and ZT was evaluated with 2-way ANOVAs and, when appropriate, differences between groups at each ZT were evaluated with unpaired t tests. Pearson correlation tests were conducted to determine the relationship between two continuous variables. Comparisons were considered significant when p < 0.05.

## Results

We and others have previously reported that serum glucocorticoid concentrations of undernourished mice are higher than controls [20,26,28]; however, it is unknown if glucocorticoid concentrations in undernourished mice exhibit diurnal variations similar to those of controls. Therefore, we measured serum corticosterone in undernourished and control mice at ZT0 and ZT12, the first hour of light and dark, respectively. Undernourished mice demonstrated serum corticosterone levels 4-6 times higher than well-nourished mice at ZT0 and ZT12 (Fig 1A). The fold change in corticosterone levels between under- and well-nourished mice was slightly greater during the light phase at ZT0, when mice are less active (Fig 1A). In summary, undernutrition elevates serum corticosterone during both the light and dark phase.

**Fig 1.**
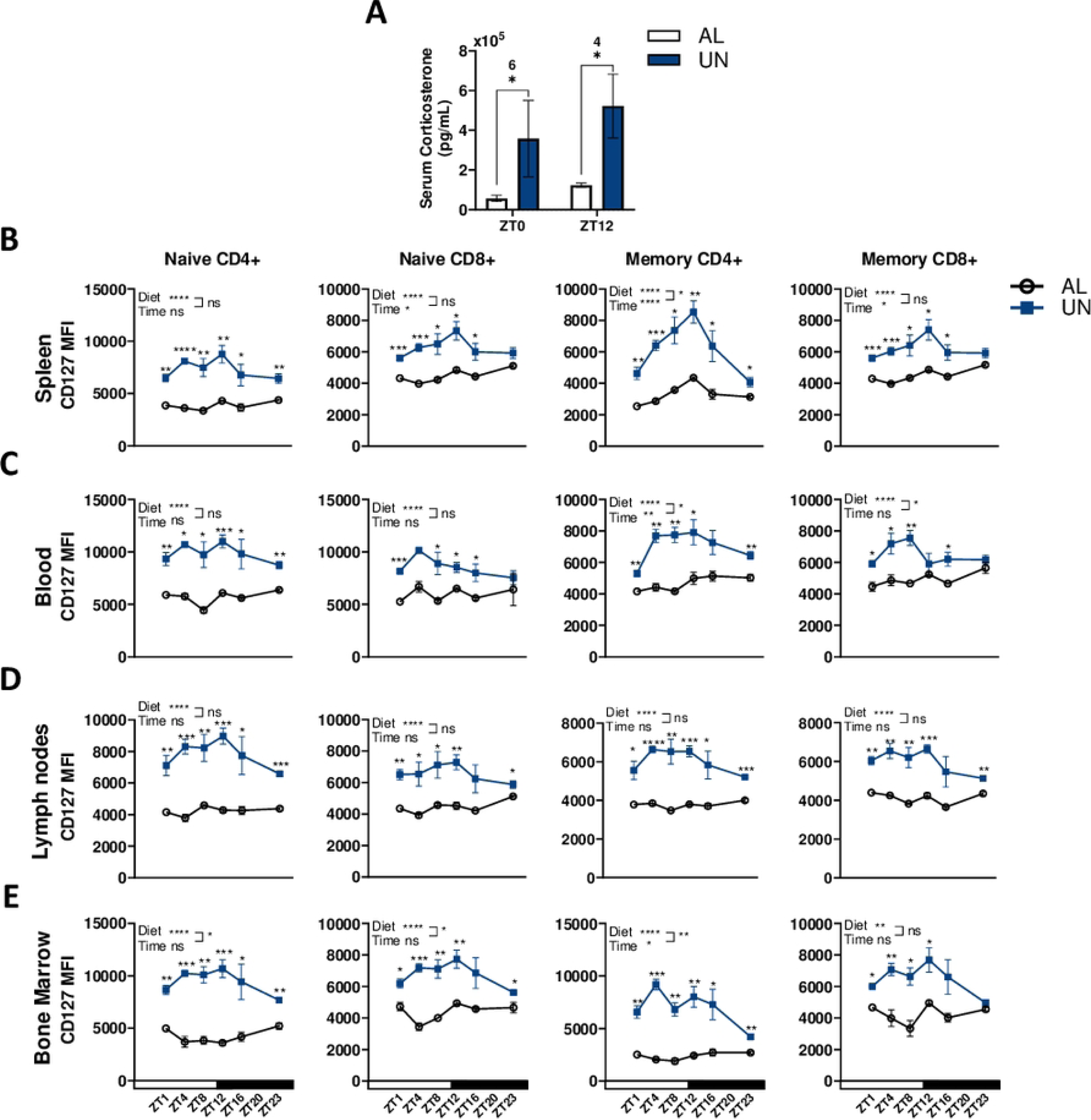
Serum corticosterone levels and CD127 expression in T cells are elevated in undernourished mice compared to well-nourished mice. Male C57BL/6J mice received a 56% reduction of standard chow (Undernourished, UN) or *ad libitum* access (AL) for 1 week. (A) Serum corticosterone levels were measured via ELISA at zeitgeber time (ZT) 0 and 12. Numbers above significance indicate the fold change in serum corticosterone levels between *AL* and UN. The median fluorescence intensity (MFI) of CD127 on naïve (CD62L^hi^) and memory (CD62L^lo^) CD4+ and CD8+ T cells isolated from the spleen (B), blood (C), lymph nodes (D), and bone marrow (E) at ZT 1, 4, 8, 12, 16 and 23 of *AL* and UN mice was evaluated by flow cytometry. n=2-4/group in 6 independent experiments conducted at 6 different ZT; by 2-way ANOVA and when appropriate by unpaired t test, ****p<0.0001, ***p<0.001, **p<0.01, *p<0.05. Each data point represents a mean ± SEM.

Glucocorticoids directly enhance CD127 expression in T cells in a diurnal fashion, which in turn is required for appropriate diurnal T cell redistribution between the blood and lymphoid organs [16]. We have previously shown that CD127 is dramatically upregulated on T cells in undernourished mice [26]. Therefore, we examined the effect of undernutrition on diurnal fluctuations in CD127 expression. We report that CD127 expression is dramatically elevated on naïve (CD62L^hi^) and memory (CD62L^lo^) CD4+ and CD8+ T cells isolated from the spleens, blood, lymph nodes, and bone marrow of undernourished mice compared to controls throughout a 24-hour period, especially during the light phase, with CD127 expression between the undernourished and controls slightly converging during the latter part of the dark phase (Fig 1B-E). In contrast to the overall increases in CD127 expression on undernourished T cells, diurnal fluctuations in expression of CD127 are similar between T cells isolated from undernourished and control mice in all tissues, except naïve CD4+ T cells isolated from the bone marrow of undernourished mice. (Fig 1B-D). In nearly all tissues isolated from both under- and well-nourished mice, peak T cell expression of CD127 occurs at or near ZT12. Interestingly, peak expression of CD127 in undernourished naïve CD4+ T cells isolated from the bone marrow is also at ZT12, although ZT12 is the trough of CD127 expression for cells isolated from well-nourished mice.

Undernourishment has been shown to reduce the total number of circulating T cells, but previous studies focused on only one timepoint, usually during the daytime, which is the inactive period for mice [20,26]. Therefore, we evaluated the total number of naïve (CD62L^hi^) and memory (CD62L^lo^) T cells at multiple ZT in undernourished and control mice to examine the influence of undernutrition on the diurnal distribution of T cells. In the spleen and blood, total numbers of naïve CD4+ and CD8+ T cells were similar in undernourished and control mice at all timepoints (Fig 2A-B). Similarly, the number of naïve T cells in the lymph nodes of undernourished and control mice were equivalent during the dark phase but during the light phase, control naïve CD8+ T cells were approximately twice as numerous as undernourished cells (Fig 2C). Interestingly, the bone marrow of undernourished mice demonstrated 2-6 times the number of T cells as well-nourished mice at nearly all ZT, with especially high numbers of naïve CD4+ and CD8+ T cells at ZT8 during the light phase (Fig 2D). Memory CD4+ and CD8+ T cells were consistently half as numerous in undernourished spleen, blood, and lymph nodes during the light phase compared to controls (Fig 2A-C). Similarly, undernourished mice demonstrated elevated numbers of memory T cells in the bone marrow compared to controls only during the light phase (Fig 2D). Given the unusual presence of naïve CD4+ and CD8+ T cells in the bone marrow of undernourished mice, we further characterized their phenotype using antibodies to CD44 and CD69 at ZT0 (Fig 2E). In the bone marrow, undernourished mice exhibited a dramatic, eight-fold increase in the number of naïve CD4+ T cells (CD62L^hi^CD44^−^ CD69^−^) and three times the number of naïve CD8+ T cells compared to well-nourished mice (Fig 2F). The bone marrow of undernourished mice also contained 2-3 times as many CD4+ and CD8+ T_CM_ cells (CD62L^hi^CD44^+^CD69^−^) than well-nourished mice (Fig 2F). There was a trend for undernourished bone marrow to harbor more CD8+ T_EM_ cells (CD62L^lo^CD44^+^CD69^−^) than well-nourished, but this was not statistically significant, while CD4+ T_EM_ cells present in similar proportions in both groups (Fig 2F). Despite the increased number of T cells in the bone marrow of undernourished mice, under- and well-nourished mice demonstrated similar total numbers of naïve and T_CM_ cells in the spleen and lymph nodes (Fig S1A-B). In the spleen, both CD4+ and CD8+ T_EM_ cell numbers were similar between both groups (Fig S1C). However, undernourished mice had half the number of CD8+ T_EM_ cells as controls in the lymph nodes (Fig S1C). Collectively, the data indicate that undernutrition significantly reduces the number of T cells in the lymph nodes, but this reduction is mostly limited to the light phase, particularly ZT8 for naïve T cells. Further, T cell numbers in the bone marrow of undernourished mice are significantly increased throughout most ZT when compared to well-nourished mice. For naïve T cells, this increase is most dramatic at ZT8 during the light phase, which is also the trough of naïve T cell residency in the lymph nodes.

**Fig 2.**
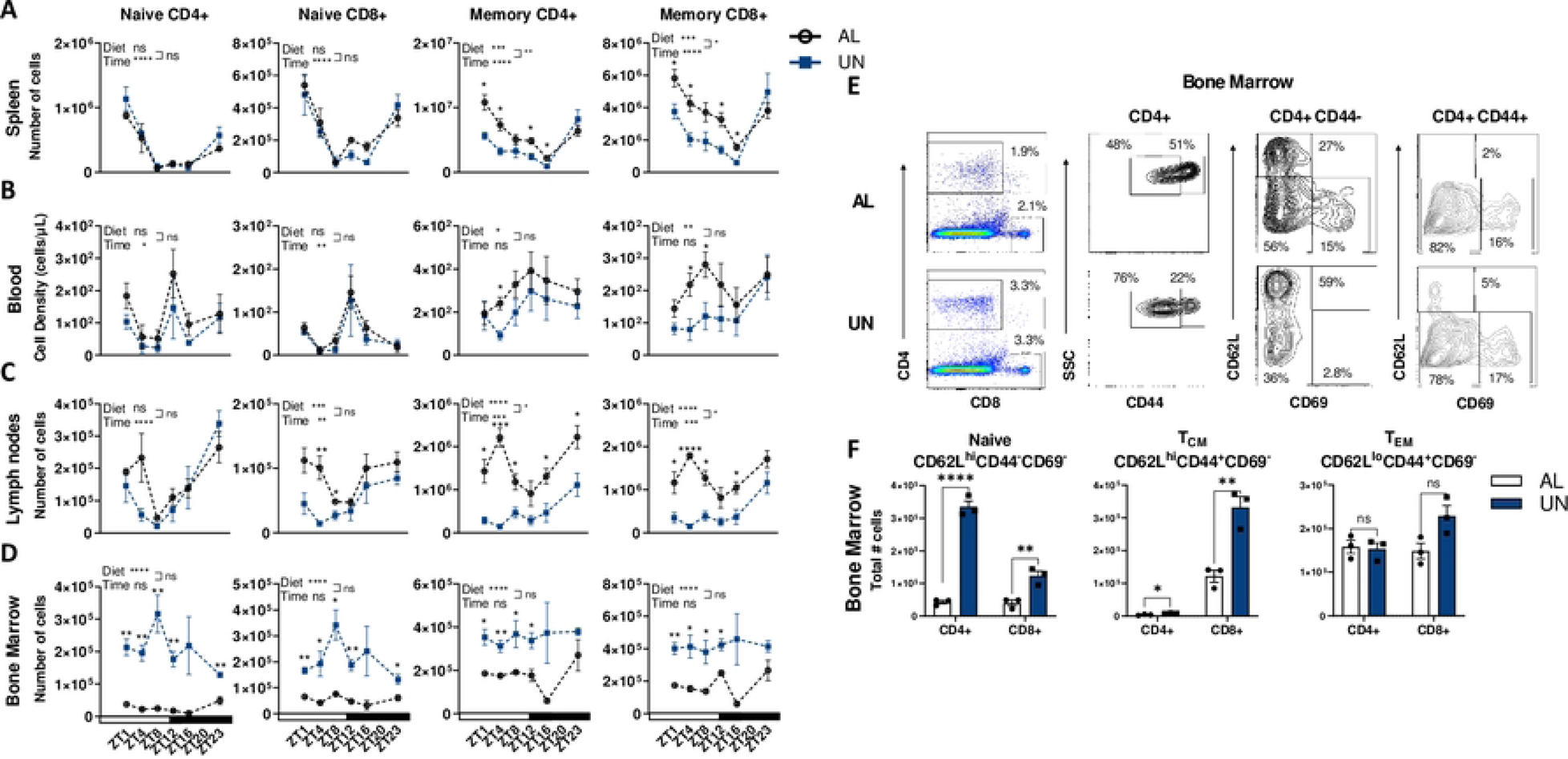
Undernourished naïve T cell numbers are increased in the bone marrow relative to well-nourished (AL), however, naïve and memory T cells favor the peripheral lymphoid organs during the dark phase in both groups. Male C57BL/6J mice received a 56% reduction of standard chow (undernourished, UN) or *ad libitum* access (AL) for 1 week. Naïve (CD62L^hi^) and memory (CD62L^lo^) CD4+ and CD8+ T cell numbers in the spleen (A), blood (B), lymph nodes (C), and bone marrow (D) were determined by flow cytometry at ZT 1, 4, 8, 12, 16, and 23. Each data point represents a mean ± SEM. n=2-3/group in 6 independent experiments conducted at 6 different ZT; by 2-way ANOVA and when appropriate by unpaired t test. The phenotype of naïve and memory T cells was additionally assessed via flow cytometry with antibodies against CD44 and CD69 at ZT0 (E). Total number of naïve, T_CM_, and T_EM_ CD4+ and CD8+ cells in the bone marrow are shown (F). n=3/group in a single independent experiment at ZT0; by unpaired t-test, ****p<0.0001, ***p<0.001, **p<0.01, *p<0.05 Graphs display means ± SEMs.

We next assessed the impact of undernutrition on T cell expression of key migration molecules to better understand what might be driving naïve T cell migration to the bone marrow. We examined CXCR4 first, since circadian changes in CXCR4 expression are controlled by GC-mediated changes in CD127 expression and CXCR4 has been associated with T cell homing to the spleen and to the bone marrow [16,29]. In undernourished mice, we found that CXCR4 expression on naïve (CD62L^hi^) CD4+ and CD8+ T cells in the spleen was modestly increased by 22% and 12%, respectively, compared to controls in both the light and dark phase (Fig 3A). Similarly, expression in the blood modestly increased by up to 20% across most ZT compared to controls (Fig S2A). In the lymph nodes, the effect of undernutrition varied by phase and T cell type. Like in the spleen, undernutrition increased CXCR4 expression on naïve CD4+ T cells, but only during the dark phase; while for naïve CD8+ T cells, undernutrition was instead associated with reduced CXCR4 expression only during the light phase (Fig 3B). Undernutrition did not dramatically affect CXCR4 expression in naïve T cells isolated from the bone marrow (Fig S2B). Undernutrition increased CXCR4 expression on memory T cells isolated from the spleen and blood but had little effect on cells isolated from the lymph nodes or bone marrow (Fig S2C-F). In summary, in the lymph nodes of undernourished mice, naïve CD4+ T cells express more CXCR4 than controls during the dark phase, a time when undernourished naïve T cell residency is equivalent to that of well-nourished mice, whereas the association is different for naïve CD8+ T cells in the lymph node. Naïve CD8+ T cells isolated from the lymph nodes of undernourished mice express less CXCR4 than controls during the light phase, a time when undernourished mice experience a deficit in naïve CD8+ T cell residency.

**Fig 3.**
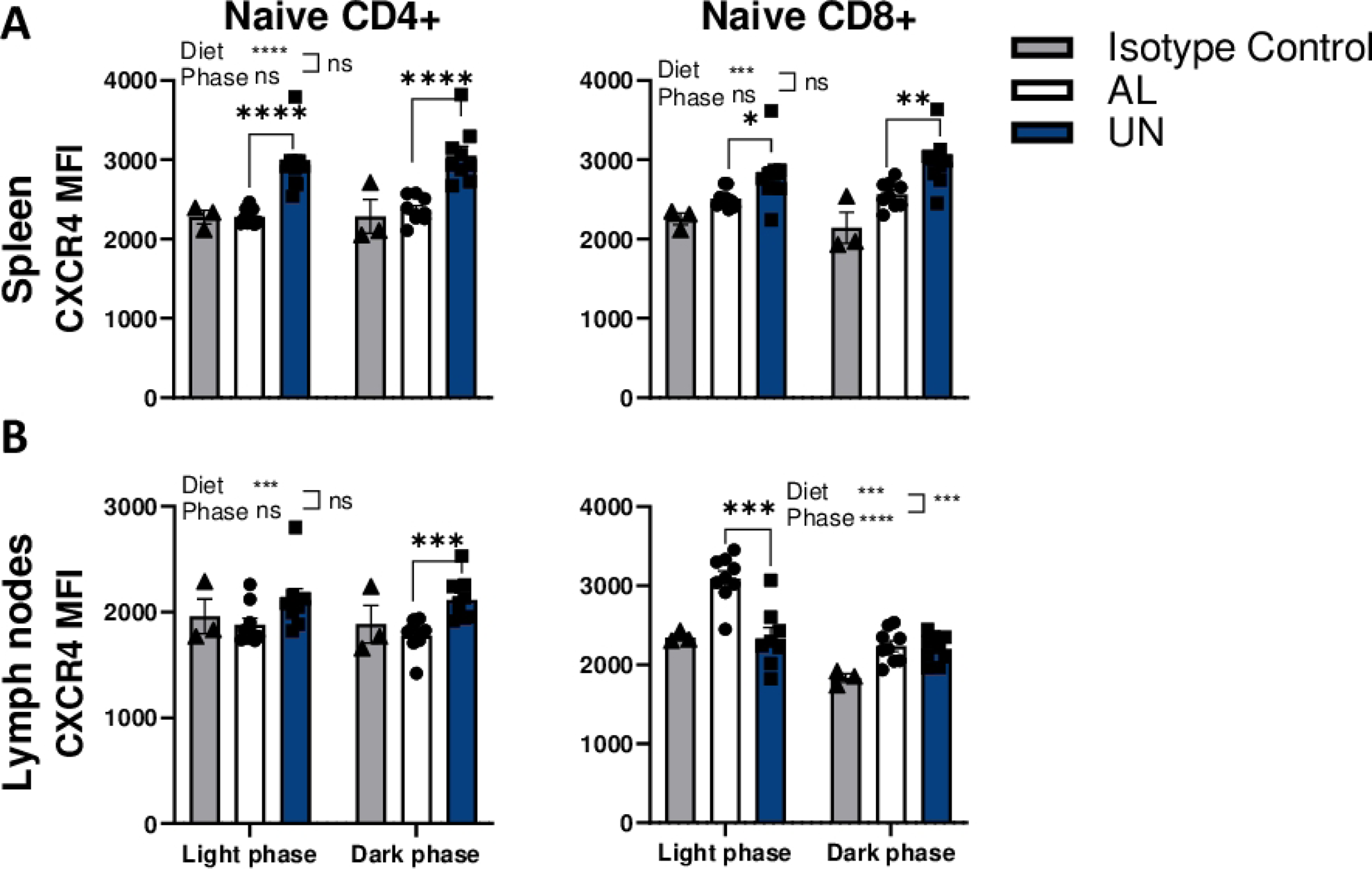
CXCR4 expression on naïve T cells is increased in the spleens of undernourished mice, while in the lymph node, undernutrition-related changes in expression are phase-dependent. Male C57BL/6J mice received 1 week of UN or *AL* treatment. The median fluorescence intensity (MFI) of CXCR4 on naïve (CD62L^hi^) CD4+ and CD8+ T cells was analyzed by flow cytometry in the spleen (A) and lymph nodes (B) during the light phase (ZT 1, 4, and 8) and the dark phase (ZT 12, 16, and 23). n=8-9/phase from 6 independent experiments conducted at 6 different ZT; by 2-way ANOVA and when appropriate by unpaired t test. ****p<0.0001, ***p<0.001, **p<0.01, *p<0.05. Each data point represents a mean ± SEM.

The rhythmic expression of CCR7 and S1P1 is associated with T cell homing and egress from the lymph nodes during the dark and light phases, respectively [22], so we examined the influence of undernutrition on their expression in T cells isolated from the lymph nodes and bone marrow. We especially focused our attention on ZT8, the height of naïve T cell occupancy in the bone marrow and near the trough of T cell occupancy of the lymph nodes. Naive CD4+ T cells in both the lymph nodes and bone marrow experienced changes in CCR7 expression during undernutrition, however in opposing directions, while expression levels on naïve CD8+ T cells were unaffected by undernutrition (Fig 4A-B). In the lymph nodes, naive CD4+ T cells from undernourished mice expressed 33% less CCR7 than their well-nourished counterparts (Fig 4A). However, naïve CD4+ T cells isolated from the bone marrow of undernourished mice expressed 30% higher levels of CCR7 compared to well-nourished mice (Fig 4B). Thus, naïve CD4+ T cells isolated from the lymph nodes of undernourished mice express less CCR7 while those from the bone marrow express more CCR7 than controls during the height of their occupation of the bone marrow. This reduction in CCR7 expression during undernutrition may serve to speed naive T cell egress from the lymph nodes [30], while increased CCR7 in bone marrow may direct the naive T cells dwelling in the bone marrow back to the lymph nodes for the active period.

**Fig 4.**
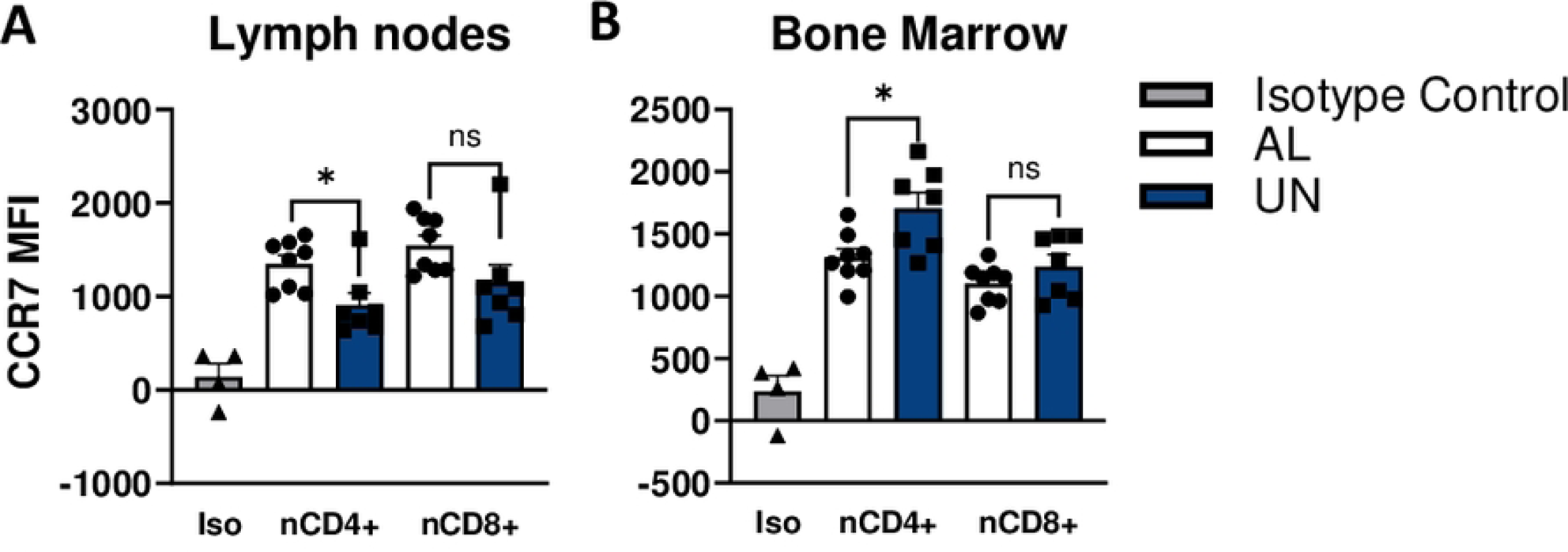
Undernutrition is associated with decreased expression of CCR7 in the lymph nodes and increased expression of CCR7 in the bone marrow on naive CD4+ T cells. Male C57BL/6J mice received 1 week of UN or *AL* treatment. The median fluorescence intensity (MFI) of CCR7 on naïve (CD44-) CD4+ and CD8+ T cells was analyzed by flow cytometry in the lymph nodes (A) and bone marrow (B) at ZT8. Graphs display means ± SEMs; n=7-8/group in two independent experiment at ZT8; by unpaired t-test, *p<0.05.

In well-nourished mice, S1P1 expression has an inverse relationship with T cell numbers, consistent with the role of S1P1 in T cell egress from lymphoid organs [31]. Hypothesizing that S1P1 might be overexpressed in T cells isolated from the lymph nodes, but under expressed in cells isolated from the bone marrow of undernourished mice, we examined S1P1 expression on T cells isolated from lymphoid tissues of under- and well-nourished mice. However, we found that S1P1 expression was mostly similar across ZT in T cells isolated from the spleen, blood, and lymph nodes of undernourished mice as compared to well-nourished controls (Figs 5A and S3A-B). On the other hand, naïve CD4+ and memory CD8+ T cells isolated from the bone marrow of well-nourished mice expressed moderately higher levels of S1P1 than cells isolated from undernourished mice, but this was limited to two ZT (Fig 5B).

**Fig 5.**
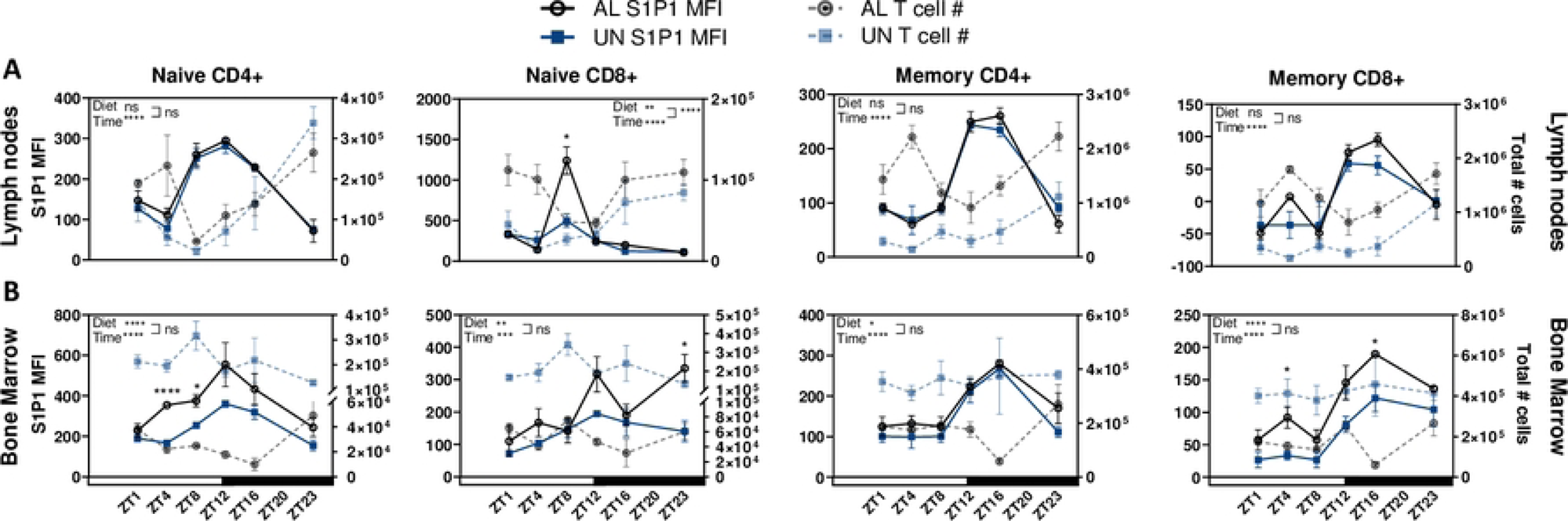
Undernourished T cells isolated from the bone marrow express less S1P1 than controls and are not negatively correlated with S1P1 expression. Male C57BL/6J mice received 1 week of UN or *AL* treatment. The median fluorescence intensity (MFI) of S1P1 on naïve (CD62L^hi^) and memory (CD62L^lo^) CD4+ and CD8+ T cells was analyzed by flow cytometry in the lymph nodes (A) and bone marrow (B) at ZT 1, 4, 8, 12, 16, and 23. n=3/group in 6 independent experiments conducted at 6 different ZT; by 2-way ANOVA and when appropriate by unpaired t test. ****p<0.0001, ***p<0.001, **p<0.01, *p<0.05. T cell numbers are indicated via dotted lines on from the right x-axis. Each data point represents a mean ± SEM.

We next considered whether undernutrition affects the relationship between S1P1 expression and T cells numbers. In the blood and spleen, both under- and well-nourished mice demonstrated the expected inverse relationship between S1P1 and the number of T cells: at ZT with high S1P1 expression in T cells, naïve (CD62L^hi^) and memory (CD62L^lo^) T cell numbers were low, and vice-versa (Fig S3). Similarly, expression of S1P1 on naïve T cells isolated from the lymph nodes of well-nourished mice demonstrated an inverse relationship with T cell numbers (Figs 5A and S4A). However, this inverse relationship was not observed during the light phase for naïve T cells isolated from the lymph nodes of undernourished mice (Figs 5A and S4B). In general, the relationship between S1P1 expression and the number of memory T cells in the lymph node was less dramatic as compared to the relationship with naïve T cells and was not observed during the light phase for memory CD8+ T cells isolated from either control or undernourished mice (Fig 5A). The number of memory CD4+ T cells isolated from the lymph nodes of well-nourished mice demonstrated an inverse relationship with S1P1 expression across ZT, but those isolated from undernourished mice only demonstrated the relationship in the dark phase (Figs 5A and S5A-B). In the bone marrow, the inverse relationship between S1P1 expression and T cell numbers was less dramatic and was mostly limited to one phase of the 24-hour period, even in control mice. The number of naïve CD4+ T cells isolated from the bone marrow of well-nourished mice demonstrated a robust inverse relationship with S1P1 expression only during the light phase; while the number of memory CD4+ T cells tended towards an inverse relationship only in the dark phase (Figs 5B, S6A-B, and S7A-B). On the other hand, the number of naïve and memory CD4+ T cells isolated from the bone marrow of undernourished mice did not exhibit an inverse relationship with S1P1 expression at any phase (Figs 5B, S6A-B, and S7A-B). Finally, no inverse relationship was detected between S1P1 expression and the number of naïve or memory CD8+ T cells isolated from the bone marrow of under- or well-nourished mice across ZT, though we did notice an unusual positive relationship with S1P1 expression and the number of naïve CD8+ T cells in the bone marrow of undernourished mice only during the light phase (Figs 5B and S8A-B). In sum, undernutrition disrupts the expected relationship between S1P1 expression and T cell numbers in the lymph nodes and bone marrow, especially during the light phase. In all cases except for naïve CD4+ T cells in the bone marrow, the loss of the inverse relationship is not associated with undernutrition-related effects on S1P1 expression.

Given the ability of CD69 to downregulate S1P1 we examined the influence of undernutrition on CD69 expression on memory T cells isolated from the lymph nodes and bone marrow. Generally, the percentages of memory (CD62L^lo^) T cells expressing CD69 fluctuated in a diurnal pattern in both under- and well-nourished mice (Fig 6A-B). In the lymph nodes, undernutrition enhanced the percentage of memory T cells expressing CD69 especially during the light phase (Fig 6A), likely reflecting an increased percentage of resident memory T cells [32]. In contrast to effects observed in the lymph nodes, a significantly smaller percentage of memory T cells isolated from the bone marrow of undernourished mice expressed CD69 as compared to controls, especially during the light phase (Fig 6B). However, undernourished T cells in the bone marrow exhibited a similar diurnal pattern of CD69 expression as control T cells (Fig 6B).

**Fig 6.**
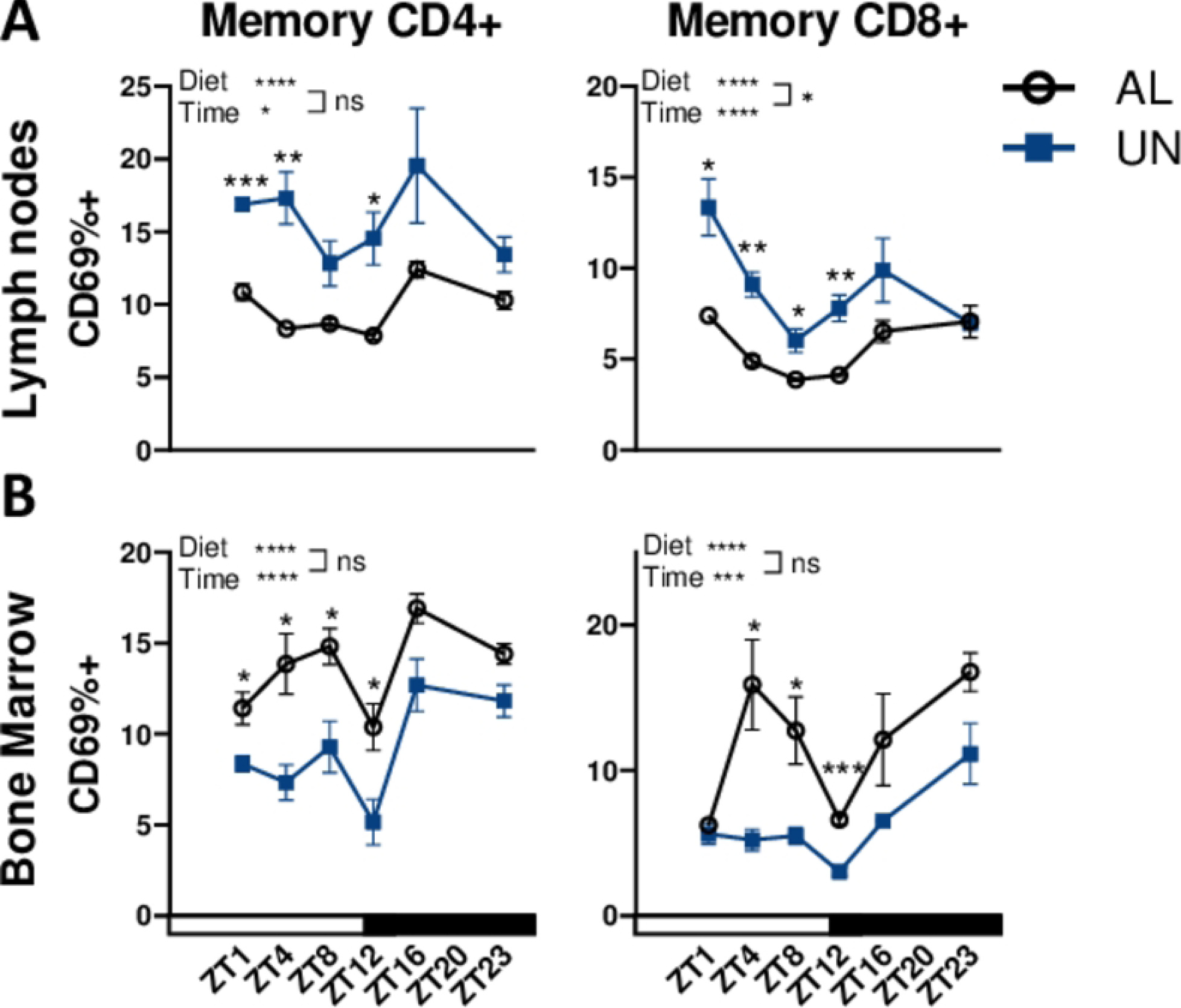
Undernutrition is associated with an increased proportion of memory T cells expressing CD69 in the lymph nodes, but a reduced proportion of CD69+ memory cells in the bone marrow, only during the light phase. Male C57BL/6J mice received 1 week of UN or *AL* treatment. The percentage of memory (CD62L^lo^) CD4+ and CD8+ T cells expressing CD69 was determined by flow cytometry in the lymph nodes (A) and bone marrow (B) at ZT 1, 4, 8, 12, 16, and 23. n=3/group in 6 independent experiments conducted at 6 different ZT; by 2-way ANOVA and when appropriate by unpaired t test. ****p<0.0001, ***p<0.001, **p<0.01, *p<0.05. Graphs display means ± SEMs.

## Discussion

In summary, undernutrition exerts the most dramatic influences on serum corticosterone levels, diurnal patterns in naïve T cell surveillance, and the expression of proteins known to control T cell migration during the light phase, when mice are less likely to encounter a pathogen (Fig 7). Indeed, undernutrition-related decreases in the number of naïve T cells in the lymph nodes are limited to the light phase (Fig 7A). Though increases in the number of naïve T cells in the bone marrow of undernourished mice are phase-independent, peak naïve T cell residency of undernourished bone marrow is also in the light phase (Fig 7B). Naïve T cells isolated from undernourished mice express higher levels of CD127 than controls across ZT, but undernutrition affects the expression of CXCR4 on CD4+ and CD8+ naïve T cells in the lymph nodes in an unexpected and phase-dependent fashion (Fig 7A). Undernourished naïve CD4+ T cells express higher levels of CXCR4 during the dark phase, when the number of naïve T cells in the lymph nodes is similar to that in control mice, while undernourished naïve CD8+ T cells express lower levels of CXCR4 during the light phase when the number of naïve T cells in the lymph nodes is deficient (Fig 7A). Naïve CD4+ T cells isolated from the lymph nodes of undernourished mice also have a lower expression of CCR7 than control mice during the light phase (Fig 7A). Moreover, naïve T cells in the lymph nodes of undernourished mice have lost the inverse relationship with S1P1 that controls naïve T cell exit (Fig 7A). Bone marrow phase-independent effects of undernutrition on naïve T cells include increases in T cell numbers and increased CD127 expression (Fig 7B). Phase-dependent effects of undernutrition in the bone marrow occur only in the light phase. During this phase, only naive CD4+ T cells lack the expected inverse relationship with S1P1, have reduced levels of S1P1, and express higher levels of CCR7 compared to controls (Fig 7B).

**Fig 7.**
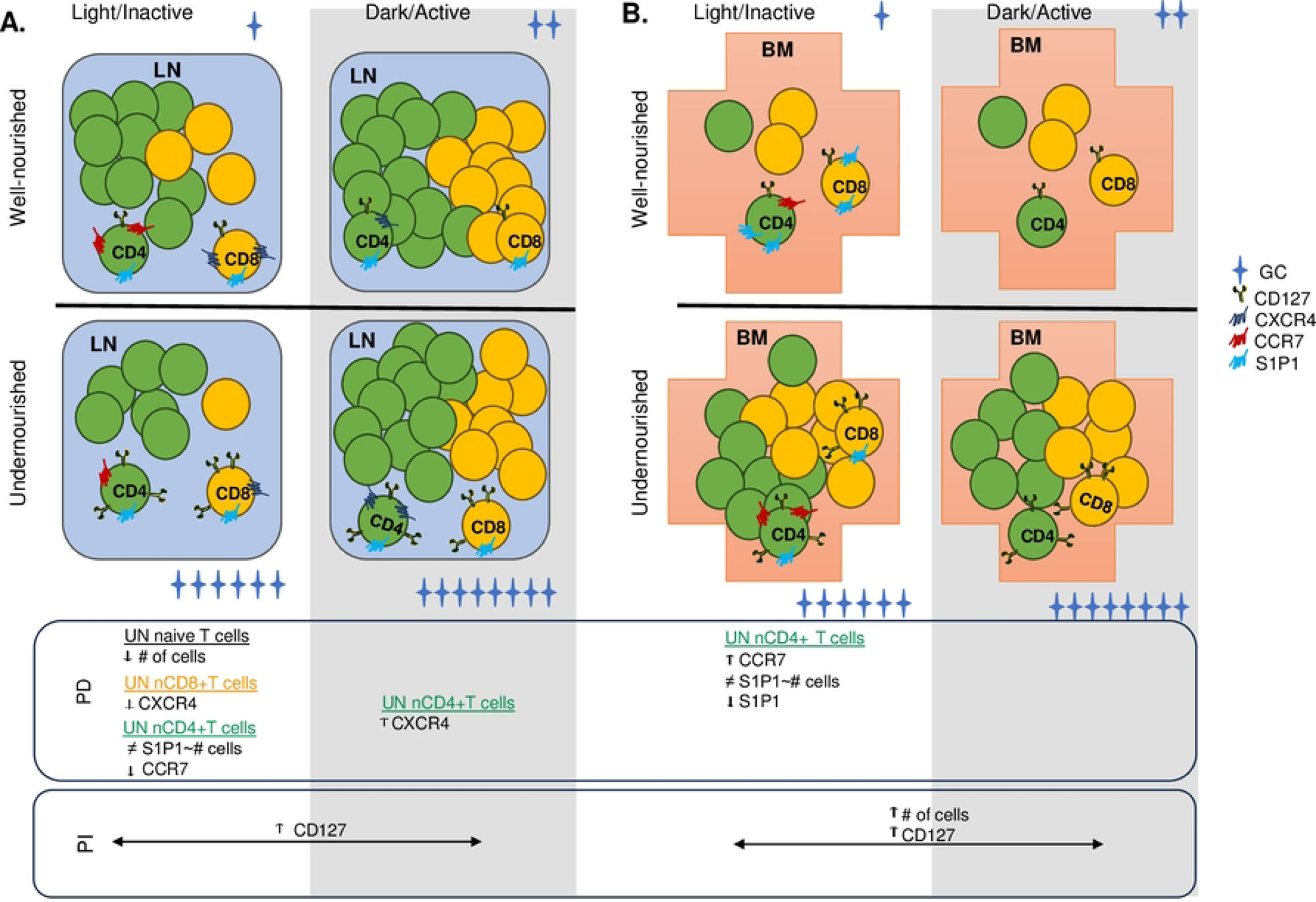
Summary of phase-dependent (PD) and phase-independent (PI) effects of undernutrition on naïve T cells in the lymph nodes and bone marrow. Undernutrition is primarily associated with PD effects on naïve T cells isolated from the lymph nodes (A) and bone marrow (B).

Our results suggest that the naïve CD4+ T cell population may be preserved in undernourished mice by sheltering in the bone marrow during the light phase. Their recirculation back to lymph nodes during the dark phase, the active period for mice, may preserve immunosurveillance when it is most necessary. Mechanistically, perhaps increased expression of CXCR4 on naïve CD4+ T cells in the lymph nodes of undernourished mice during the dark phase sends the cells to the bone marrow for the light phase and increased CCR7 expression on the same cells in the bone marrow during the light phase directs them back to the lymph nodes for the dark phase. Given that all T cell subsets isolated from undernourished mice upregulate CD127, we do not believe, as originally hypothesized, that CD127 is responsible for the observed phase and tissue specific changes in CXCR4 expression in naïve T cells. In well-nourished mice, mTORC2-mediated suppression of CXCR4 is critical for preventing naïve T cells from migrating to the bone marrow [33]. Perhaps undernutrition releases naïve CD4+ T cells from this ‘brake’ on bone marrow homing. Additionally, GCs can enhance T cell responsiveness to the CXCR4 ligand, CXCL12, in a manner that does not involve up-regulation of CXCR4 nor the transcription-modifying activity of GR [34]. Perhaps the highly elevated levels of serum corticosterone reported here enhance naïve T cell responsiveness to CXCL12 and results in their migration to the bone marrow. Naïve CD4+ T cell disengagement from S1P1-related egress signals may further contribute to increased retention of T cells within each compartment during the appropriate phase. It is less clear what might be driving increases in naïve CD8+ T cell migration to the bone marrow during undernutrition, especially since these cells downregulate CXCR4 expression while in the lymph node during the light phase.

During undernutrition, memory T cells are reduced in number in the lymph nodes and other secondary lymphoid organs across ZT but are more dramatically deficient in the lymph nodes during the light phase (Fig 8A). In addition, undernourished memory cells express higher levels of CD127 than well-nourished controls across ZT (Fig 8A). Light phase specific undernutrition-related changes in the lymph nodes include increases in the proportion of memory T cells expressing CD69 compared to controls and the lack of an inverse relationship between S1P1 expression and the number of memory CD4+ T cells (Fig 8A). Memory T cells isolated from the bone marrow of undernourished mice were greater in number, expressed more CD127, and expressed less S1P1 than cells isolated from controls across ZT (Fig 8B). Memory CD4+ T cells isolated from the bone marrow of undernourished mice did not exhibit an inverse relationship with S1P1 expression during the dark phase, even though control cells did (Fig 8B). Finally, during the light phase, a reduced proportion of memory T cells isolated from the bone marrow of undernourished mice expressed CD69 compared to controls (Fig 8B). It is additionally worth mentioning that undernourished memory T cells isolated from the spleen expressed high levels of CXCR4 across ZT compared to controls. Thus, our results are consistent with another study demonstrating that memory T cells are preserved in the bone marrow during dietary restriction via a mechanism involving the CXCR4:CXCL12 and S1P1:S1P signaling axes [20].

**Fig 8.**
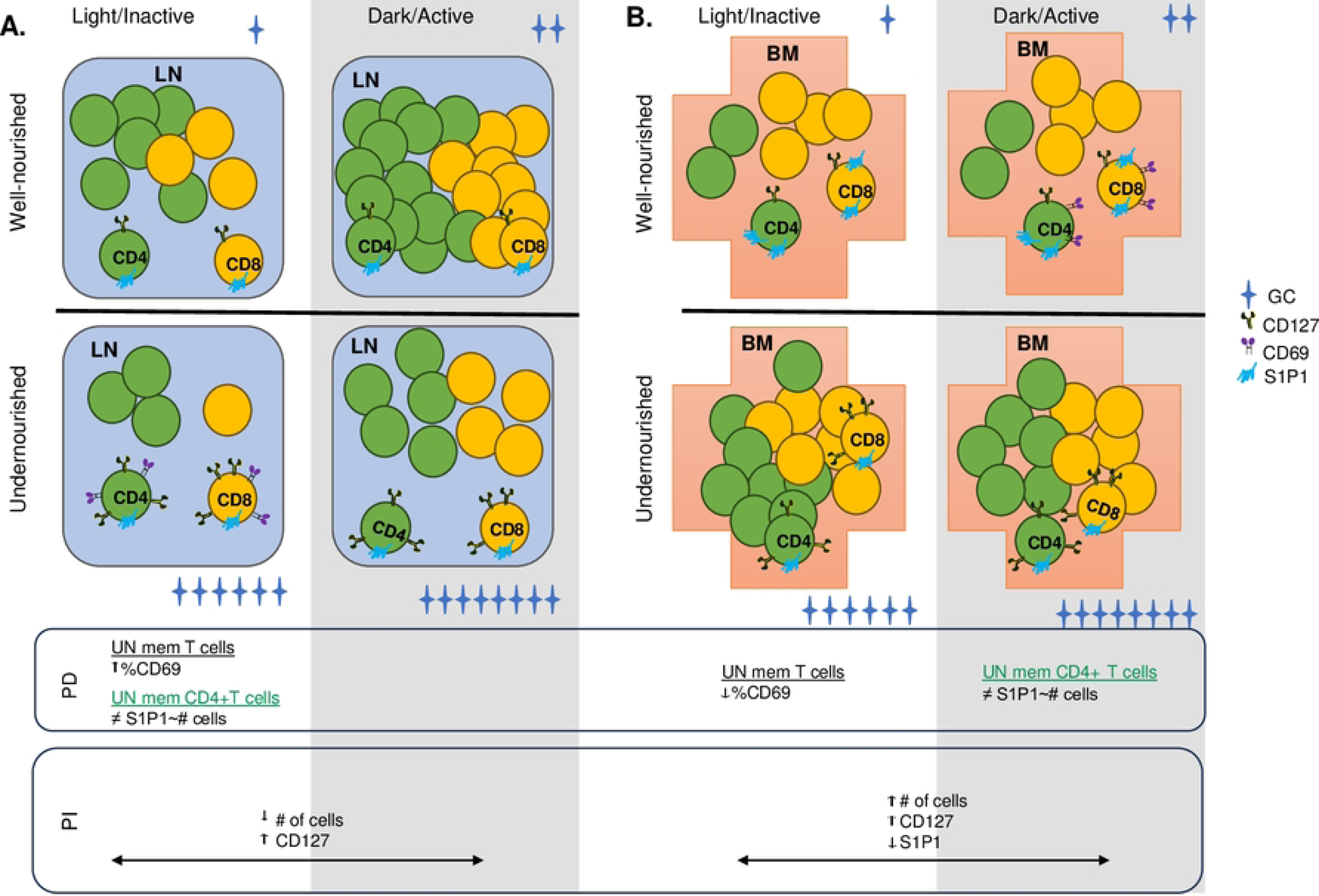
Summary of phase-dependent (PD) and phase-independent (PI) effects of undernutrition on memory T cells in the lymph nodes and bone marrow. Undernutrition is primarily associated with PI effects on memory T cells isolated from both the lymph nodes (A) and bone marrow (B).

Circadian rhythmicity of T cell trafficking affects the efficiency of immunosurveillance as well as the strength and type of immune response generated at a given time of day. The T cell repertoire consists of anywhere between 25 million to 1 billion T cells, of which, only about 100 share T cell receptors (TCR) with a similar specificity to peptides derived from any single pathogen. Yet, most antigen-specific T cells are activated only three days after a new antigen is detected by the innate immune system [14]. This is possible, at least in part, because dendritic cells and T cells share the same circadian trafficking patterns to secondary lymphoid organs [35]. Moreover, the strength and type of the immune response mounted is time of day dependent for a variety of reasons including the circadian trafficking patterns of immune cells [36]. For example, vaccination in humans with the Sinopharm protein-based inactivated SARS-CoV-2 vaccine was more effective when administered at the start of the active period rather than the end [37]. Prioritizing immune cell communication in lymph nodes during the active period makes sense for at least two reasons—the active period is likely to be when feeding activity occurs and is also the time when an organism is most likely to encounter an infectious agent. In agreement with others [16,22], we report here that T cell residency in the lymph nodes of mice peak during the active period. However, while other studies have indicated that total number of T cells in the lymph node peaks at the onset of the active period, our results indicate a gradual rise throughout the active period [16,22]. Two potentially important differences in our study design may account for this slight shift—the time at which the chow was replenished and the housing conditions. Although the control mice in our experiments were fed *ad libitum*, they were given fresh food at approximately ZT12, to best mimic their natural feeding time. It is possible that the timing of food introduction could affect feeding behavior, and feeding activity is a known zeitgeber, at least at the cellular level. Secondly, in order to accurately measure how much chow individual mice consumed, all experimental mice were singly housed and housing conditions could affect glucocorticoid levels, which could in turn influence circadian rhythms [38,39].

The dramatic increase in the total number of naïve and memory T cells in the bone marrow of undernourished mice is striking and prompts consideration of what adaptive benefits may result. One possibility is that T cell presence in the bone marrow influences hematopoiesis to prioritize the production of specific cell types needed during caloric and nutrient deficiencies. Supportively, activated T cells coordinate peripheral demand for particular immune cells with their development in the bone marrow during infection [40]. In the absence of infection, T cell deficient mice exhibit defective myelopoiesis and erythropoiesis, and the restoration of CD4+ T cells rescues myelopoiesis, but not erythropoiesis. Myelopoiesis was only restored when the CD4+ T cells were activated in an antigen-specific manner [24]. Collins et al. noted the dramatic increase in erythropoiesis during dietary restriction [20] and we also noted the bone marrow of undernourished mice was a darker red compared to bone marrow from well-nourished mice (unpublished observation). In both studies, the number of T cells in the bone marrow was elevated. Together these observations support the hypothesis that T cell presence in the bone marrow enhances erythropoiesis but does not reveal how or why such a mechanism may have evolved. During food deprivation, organisms exhibit increased locomotor activity, which is likely an adaptation that enables foraging behavior [41]. Perhaps T cell-mediated enhanced erythropoiesis during undernutrition enables greater physical activity, even in the face of reduced nutrition and caloric intake, for more effective foraging.

In addition to the potential influence of T cells on hematopoiesis, it is possible that migration of T cells to the bone marrow during undernutrition serves to preserve T cell immunity. Indeed, Collins et al. demonstrated that CD8+ central memory T cells in the bone marrow of dietary restricted mice exhibited a transcriptional profile consistent with conserving energy and that their migration to the bone marrow was associated with improved memory cell function [20]. The association between time spent in the bone marrow and the strength of the memory response was also revealed in a different study utilizing CD69 knockout mice [25]. Effector CD4+ T cells deficient for CD69 failed to migrate to or persist in the bone marrow and were also unable to generate long-lived memory cells [25]. Perhaps the increased percentage of T cells expressing CD69 in the lymph nodes of undernourished mice reported here contributes to their migration to and retention in the bone marrow. In any case, it will be interesting to ascertain the impact of increased bone marrow residency during undernutrition on naïve T cell function.

GCs exhibit diurnal fluctuations in the serum [12], and T cell sensitivity to GCs is required for the diurnal nature of T cell surveillance and optimal immunity during the active period [15,16]. In the current study, undernutrition led to an increase in circulating GCs and the migration of naïve T cells into the bone marrow. Thus, it seems possible that the increase in circulating GCs is responsible for the accumulation of T cells in the bone marrow during undernutrition. Supportively, exposure to natural or synthetic glucocorticoids in multiple mammalian models led to the decline of T and B cells in the periphery and an accumulation of T cells in the bone marrow [42,43]; and, increased serum GCs were associated with memory CD8+ T cell trafficking into the bone marrow during dietary restriction [20]. Notably, even though memory CD8+ T cells express the glucocorticoid receptor, migration to the bone marrow during dietary restriction did not require its expression [20]. It will be important to determine if naïve T cell sensitivity to GC is required for undernutrition-mediated migration of naïve T cells into the bone marrow.

Under normal conditions, the number of T cells in lymphoid organs is inversely proportional to S1P1 expression on T cells [44]. However, we report here that this inverse relationship is not observed for naïve or memory CD4+ T cells in the lymph nodes during the light phase. The upregulation of CD69 by undernourished memory CD4+ T cells during the light phase may sequester these cells in the lymph nodes by forming a complex that initiates the downmodulation and inhibition of the S1P1 receptor. This could potentially alter T cell trafficking behavior from the lymph nodes [45]. In addition, naïve and memory CD4+ T cell numbers were not in an inverse relationship with S1P1 in the bone marrow during the light and dark phases, respectively. Unlike the light phase in the lymph nodes for memory T cells, the dark phase in the bone marrow was not accompanied by increased CD69 expression. Despite lacking a clear interaction with CD69, the role of S1P1 and its sphingolipid S1P may serve another purpose. Dietary restriction increases erythropoiesis [20], and erythrocytes appear to be the main source of S1P within plasma [46,47]. Perhaps the increased numbers of erythrocytes in the bone marrow during undernutrition eliminates the S1P gradient that may normally attract T cells into the blood. Another possibility is that S1P promotes naïve T cell survival by enhancing mitochondrial function independent of the S1P1 receptor’s role in T cell egress, which has been shown in lymphoid tissue [48].

In summary, we report that undernutrition dramatically increases T cell occupancy of the bone marrow throughout the entire day and especially during the light phase, with diminished T cell occupancy of the lymph nodes also primarily occurring during the light phase. Undernutrition-related changes in T cell expression of key migration proteins are also mostly limited to the light phase. Overall, our results are in line with the hypothesis that undernutrition-mediated naïve CD4+ T cell upregulation of CXCR4 in the lymph nodes during the dark phase enables their migration to the bone marrow where they remain for the light phase. Once there, CCR7 is upregulated, sending them back to the lymph node, thereby preserving immunosurveillance during the dark phase. Alterations in the S1P1-S1P signaling axis may additionally contribute to the distinct circadian rhythm associated with undernutrition. Undernutrition-related increases in T cell residency of the bone marrow throughout the light/dark cycle likely serve to preserve T cell numbers until nutrition is restored, though further studies examining the impact of bone marrow residency on naïve T cell function will be informative.

## Acknowledgements

We thank Jacob Hanes, Kimberly Cox, and Kate Laws for their careful and thoughtful feedback on the manuscript. We also thank Olivia Adams, Rithanya Saravanan, and Aja Washington for their technical assistance.

## Supporting information

**S1 Fig. The number of T_EM_ cells in the lymph nodes are diminished during undernutrition.** Total numbers of naïve (A), TCM (B), and TEM (C) CD4+ and CD8+ cells are shown in spleen and lymph nodes. n=2-3/group; by unpaired t-test, **p<0.01, *p<0.05. Graphs display means ± SEMs.

**S2 Fig. During undernutrition, CXCR4 expression is increased on naïve and memory T cells in the blood, and on naïve CD4+ T cells in the bone marrow.** Male C57BL/6J mice received a 56% reduction of standard chow (UN) or *ad libitum* access (AL) for 1 week. The median fluorescence intensity (MFI) of CXCR4 on naïve (CD62L^hi^) (A-B) and memory (CD62L^lo^) (C-F) CD4+ and CD8+ T cells in the spleen, blood, lymph nodes, and bone marrow was analyzed by flow cytometry at ZT 1, 4, 8, 12, 16, and 23. n=2-3/group/ZT; by 2-way ANOVA and when appropriate unpaired t tests, ****p<0.0001, ***p<0.001, **p<0.01, *p<0.05. Each data point represents a mean ± SEM.

**S3 Fig. T cell numbers in the spleen and blood generally are inversely correlated to S1P1 expression in both under- and well-nourished mice.** Male C57BL/6J mice received 1 week of UN or *AL* treatment. The median fluorescence intensity (MFI) of S1P1 in the spleen (A) and blood (B) on naïve (CD62L^hi^) and memory (CD62L^lo^) CD4+ and CD8+ T cell was analyzed by flow cytometry at ZT 1, 4, 8, 12, 16, and 23. n=2-3/group/ZT; by 2-way ANOVA and when appropriate unpaired t tests, ****p<0.0001, ***p<0.001, **p<0.01, *p<0.05. T cell numbers are indicated via dotted lines on from the right x-axis. Each data point represents a mean ± SEM.

**S4 Fig. Naïve T cell numbers in the lymph nodes of undernourished mice lack a negative correlation with S1P1 expression during the light phase.** The MFI of S1P1 on naïve T cells isolated from the lymph nodes of *ad libitum*-fed (AL) or undernourished (UN) mice and the total number of T cells at each ZT were plotted against each other and tested for significance using a Pearson correlation analysis. Each dot represents a single mouse. The r and p values from the correlation test are shown.

**S5 Fig. Memory CD4+ T cell numbers in the lymph nodes of undernourished mice lack a negative correlation with S1P1 expression during the light phase.** The MFI of S1P1 on memory CD4+ T cells isolated from the lymph nodes of *ad libitum*-fed (AL) or undernourished (UN) mice and the total number of T cells at each ZT were plotted against each other and tested for significance using a Pearson correlation analysis. Each dot represents a single mouse. The r and p values from the correlation test are shown.

**S6 Fig. The number of control, but not undernourished, naïve CD4+ T cells in the bone marrow demonstrate an inverse relationship with S1P1 expression only during the light phase.** The MFI of S1P1 on naïve CD4+ T cells isolated from the bone marrow of *ad libitum*-fed (AL) or undernourished (UN) mice and the total number of T cells at each ZT were plotted against each other and tested for significance using a Pearson correlation analysis. Each dot represents a single mouse. The r and p values from the correlation test are shown.

**S7 Fig. The number of control, but not undernourished, memory CD4+ T cells in the bone marrow trend towards an inverse relationship with S1P1 expression only during the dark phase.** The MFI of S1P1 on memory CD4+ T cells isolated from the bone marrow of *ad libitum*-fed (AL) or undernourished (UN) mice and the total number of T cells at each ZT were plotted against each other and tested for significance using a Pearson correlation analysis. Each dot represents a single mouse. The r and p values from the correlation test are shown.

**S8 Fig. Naïve CD8+ T cell numbers in the bone marrow are positively correlated with S1P1 expression in undernourished mice during only the light phase.** The MFI of S1P1 on naive CD8+ T cells isolated from the bone marrow of *ad libitum*-fed (AL) or undernourished (UN) mice and the total number of T cells at each ZT were plotted against each other and tested for significance using a Pearson correlation analysis. Each dot represents a single mouse. The r and p values from the correlation test are shown.

## Notes

### Competing Interest Statement

The authors have declared no competing interest.

